# Structural basis of epitope selectivity and potent protection from malaria by PfCSP antibody L9

**DOI:** 10.1101/2022.10.07.511358

**Authors:** Gregory M. Martin, Monica L. Fernández Quintero, Wen-Hsin Lee, Tossapol Pholcharee, Lisa Eshun-Wilson, Klaus R. Liedl, Marie Pancera, Robert A. Seder, Ian A. Wilson, Andrew B. Ward

## Abstract

A primary objective in malaria vaccine design is the generation of high-quality antibody responses against the circumsporozoite protein of the malaria parasite, *Plasmodium falciparum* (PfCSP). To enable rational antigen design, we solved a cryo-EM structure of the highly potent anti-PfCSP antibody L9 in complex with recombinant PfCSP. We found that L9 Fab binds multivalently to the CSP minor (NPNV) repeats, which is stabilized by a novel set of affinity-matured homotypic, antibody-antibody contacts. Molecular dynamics simulations revealed a critical role of the L9 light chain in integrity of the homotypic interface, which likely impacts CSP affinity and protective efficacy. These findings reveal the molecular mechanism of the unique NPNV selectivity of L9 and emphasize the importance of anti-homotypic affinity maturation in protective immunity against *P. falciparum*.

**One sentence summary:** The L9 light chain is crucial for potency by conferring multivalent, high affinity binding to the NPNV minor repeats of PfCSP.

## Main Text

Malaria remains one of the world’s deadliest infectious diseases, and in 2021 was responsible for 241 million clinical infections and 627,000 deaths worldwide (*1*), primarily among young children in sub-Saharan Africa. RTS,S/AS01B (RTS,S), the only approved malaria vaccine, is only partially effective, providing ~30% protection from clinical infection after four years in children aged 5-17 months (*2*, *3*). Thus new tools, like next-generation vaccines and highly potent monoclonal antibodies (mAbs), the latter of which can provide prolonged, sterilizing immunity, are needed for prevention and elimination of malaria.

PfCSP, the primary surface antigen of *P. falciparum* sporozoites, is a major target for vaccines and mAbs as it is both highly conserved and critical for the initiation of malaria infection. PfCSP contains an immunodominant central repeat region composed of repeating four-amino-acid units, structurally defined as DPNA, NPNV, and NPNA (*4*–*11*). These roughly define the junctional, minor repeat, and major repeat epitopes, respectively. Each epitope can generate potent antibodies that prevent malaria infection in animal models (*12*–*14*), with the junctional mAb cis43LS demonstrating high-level protection against controlled human malaria infection (CHMI) in humans (*15*). Recently, we identified the minor-repeat-specific mAb L9 as one of the most potent anti-PfCSP mAbs isolated to date (*16*), which can also confer high-level sterilizing immunity against CHMI in humans (*17*). Like many of the most potent NPNA-specific mAbs, L9 is also encoded by the *IGHV3-33/IGKV1-5* heavy/light chain gene combination. However, L9 is highly specific for the NPNV (minor) repeats and relies on critical contributions from the light chain for both NPNV selectivity and high potency (*6*).

To understand the molecular basis of these unique functional properties, we solved a 3.36 Å cryoEM structure of L9 Fab in complex with a recombinant PfCSP construct, rsCSP, which contains the full N-terminal, junctional, and C-terminal regions, and about half the number of NPNA repeats as the 3D7 reference strain (Fig. 1, fig. S1, table S1). In the cryo-EM map, we observe three tightly packed Fabs bound to a central rsCSP, with each Fab simultaneously interacting with the peptide and the adjacent Fab via homotypic interactions (*8*, *11*, *18*, *19*). In general, the complex is homogeneous and the density is well-resolved for each L9 variable region (Fv) as well as the rsCSP peptide (Fig. 1B). The structure of rsCSP, built *de novo* based on the EM density, consists solely of the minor repeat region (Fig. 1F). The modeled antigen sequence comprises 26 residues encompassing three complete NPNV and DPNA repeats, i.e. NA(NPNVDPNA)_3_; there is no additional density observed that would correspond to N-terminal, C-terminal, or major repeat regions. The L9 Fab and peptide cryo-EM structures correspond well with our recent X-ray structures of two chimeric precursors of L9 (L9_K_/F10_H_ and F10_K_/L9_H_) in complex with a short minor repeat peptide (NA<u>NPNV</u>DP) (*6*) (fig. S2). Relative to the L9 cryo-EM structure, RMSD values for both chimeric Fvs are ~0.5Å, and ~0.1Å over the NPNV peptide.

**Figure 1.**
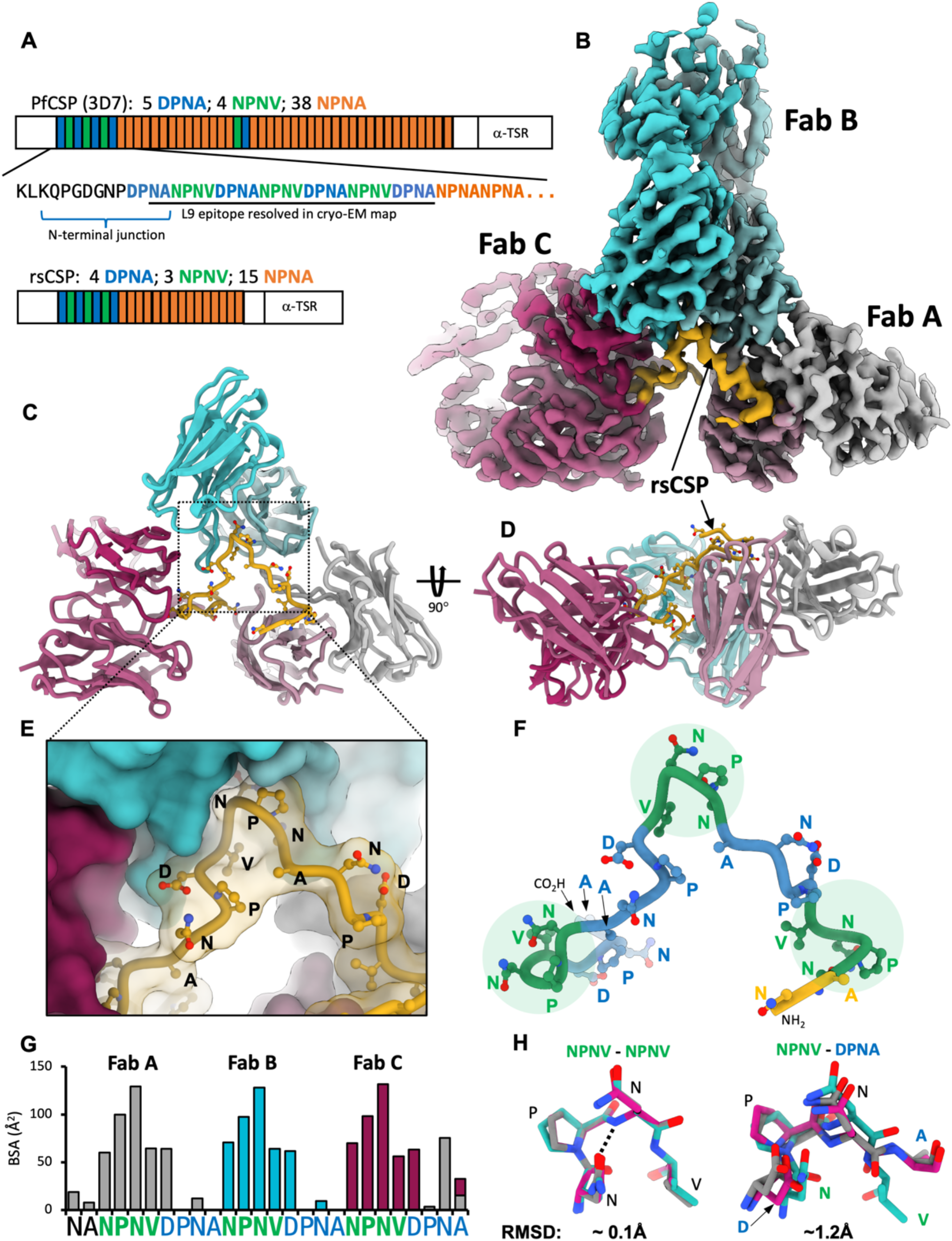
Cryo-EM structure of the L9 Fab-rsCSP complex. **(A)** Schematic of protein sequence of full-length PfCSP and rsCSP (recombinant). Each box corresponds to a single repeat. The minor repeat region is in blue and green. **(B)** Cryo-EM map of L9-rsCSP at 3.36Å. **(C)** Ribbon diagram of the atomic model; only the Fab variable region (Fv) was built into the density. **(D)** Rotated view of (C). **(E)** Zoomed-in view of (C), shown in a surface representation. **(F)** Model of the minor repeat peptide, colored as in (A). NPNV type-1 β-turns are highlighted with a green circle. **(G)** Buried surface area on rsCSP, color-coded to the Fab with which each rsCSP residue interacts. **(H)** Alignment of the three NPNV motifs (left), or the three DPNA motifs aligned to the central NPNV motif (right).

In the cryo-EM structure, each L9 Fab primarily engages a single NPNV repeat, while the DPNA repeats are largely unbound and serve as a linker between each NPNV (Fig. 1C and E). Thus, the full epitope bound by a single L9 Fab is NPNVD (Fig. 1G). Each NPNV motif adopts a type 1 β-turn, which is frequently observed for DPNA and NPNA motifs bound to anti-PfCSP antibodies from a variety of heavy chain lineages (*4*, *8*, *20*). The DPNA repeats in the L9 structure, however, are more extended and lack clear secondary structure elements (Fig. 1,F and H). The L9 epitope is centered on the NPNV type 1 β-turn, which resides in a deep, central pocket on the Fab formed primarily from CDRL1, CDRL3, and CDRH3, with smaller contributions from CDRH1 and H2 (Fig. 2A). Interestingly, overall buried surface area (BSA) on L9 is slightly biased toward the light chain (LC; L9_K_) (fig. S3, A and B). Of the 550Å^2^ total BSA on a single L9 Fab, L9_K_ contributes 294 Å^2^ (53.5%), while the heavy chain (HC; L9_H_) contributes 256Å^2^ (46.5%), indicating a critical role of L9^K^ in PfCSP binding.

**Figure 2.**
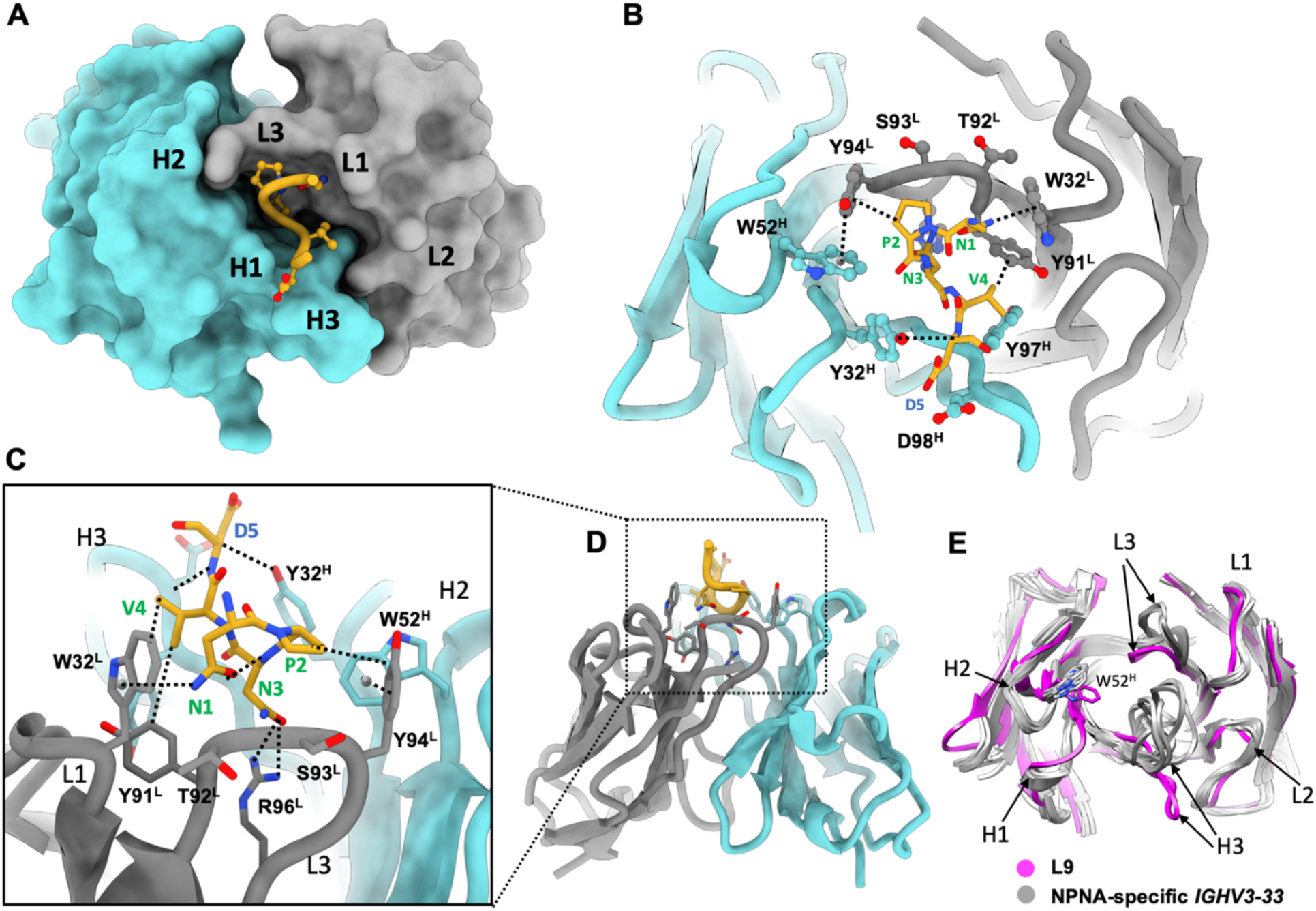
The L9 PfCSP epitope comprises NPNVD. **(A)** Surface representation of L9 Fab, with central NPNVD shown in gold. CDR loops are specified. **(B)** Structural details of CSP binding pocket. Key interactions are highlighted with dashed lines. **(C)** Rotated view of (B), zoomed in from (D). **(D)** Rotated view of (A), shown in ribbon diagram. **(E)** Alignment of L9 Fab (magenta) with a panel of NPNA-specific *IGHV3-33* Fabs; sequences in fig. S5.

As frequently observed in anti-NPNA major repeat mAbs, many direct antigen contacts are with germline-encoded aromatic residues, which in L9 create a hydrophobic cage surrounding the NPNV motif (Fig. 2B, table S2). In particular, W32^L^ in CDRL1 stacks closely against the N-terminal Asn of the NPNV motif (N1) forming a CH-π bond, while Y94^L^ in CDRL3 engages the repeat Pro (P2) (Fig. 2C, fig. S3, C and D). L9 also utilizes the strictly conserved *IGHV3-33* germline residue W52^H^ in CDRH2, which in all structures of *IGHV3-33* mAbs solved to date forms a critical CH-π interaction with P6 of the second NPNA repeat in the NPNA_2_ epitope (*7*, *8*, *11*, *20*). However, in L9, this role is assumed by Y94^L^, and W52^H^ principally acts to stabilize the Y94^L^:P2 interaction through a π-π stacking interaction with the Y94^L^ side chain (Fig. S3D-F).

This paratope structure is distinct from most other *IGHV3-33* mAbs targeting both major and minor repeats. In L9, a repositioning of the HC and LC CDR3 loops, along with a rearrangement of W52^H^ and CDRH2, creates a compact, central CSP binding pocket bounded by each of the HC and LC CDRs (Fig. 2B). A somatically mutated residue, R96^L^ in CDRL3, is found at the base of the pocket and creates a highly basic cavity (Fig. S3G). This basic binding pocket is nearly fully occupied by the N3 side chain, which forms key H-bonds with R96^L^ (Fig. 2C), while V4 occupies a hydrophobic cavity at the interface of CDRL1, L3, and CDRH3 (fig. S3C). With this unique CDR conformation, L9 appears optimally disposed to bind the bulkier minor repeat residue V4, which is the only difference between the NPNA and NPNV epitopes.

Another unique property of L9 is the ability to “crosslink” two NPNV motifs within the minor repeat region of PfCSP, which improves binding affinity (*6*). Our cryo-EM structure reveals that L9 achieves this through multivalent Fab binding to sequential NPNV repeats stabilized by an extensive antibody-antibody, or homotypic, interface between adjacent Fabs (Fig. 3A). Homotypic interactions have now been identified in several anti-NPNA mAbs and appear to be a characteristic feature of the *IGHV3-33* antibody family (*8*, *11*, *18*–*20*). Importantly, we demonstrate L9 as the first non-NPNA targeting anti-PfCSP mAb to also utilize homotypic interactions, suggesting that both the major and minor PfCSP repeats can facilitate their development.

**Figure 3.**
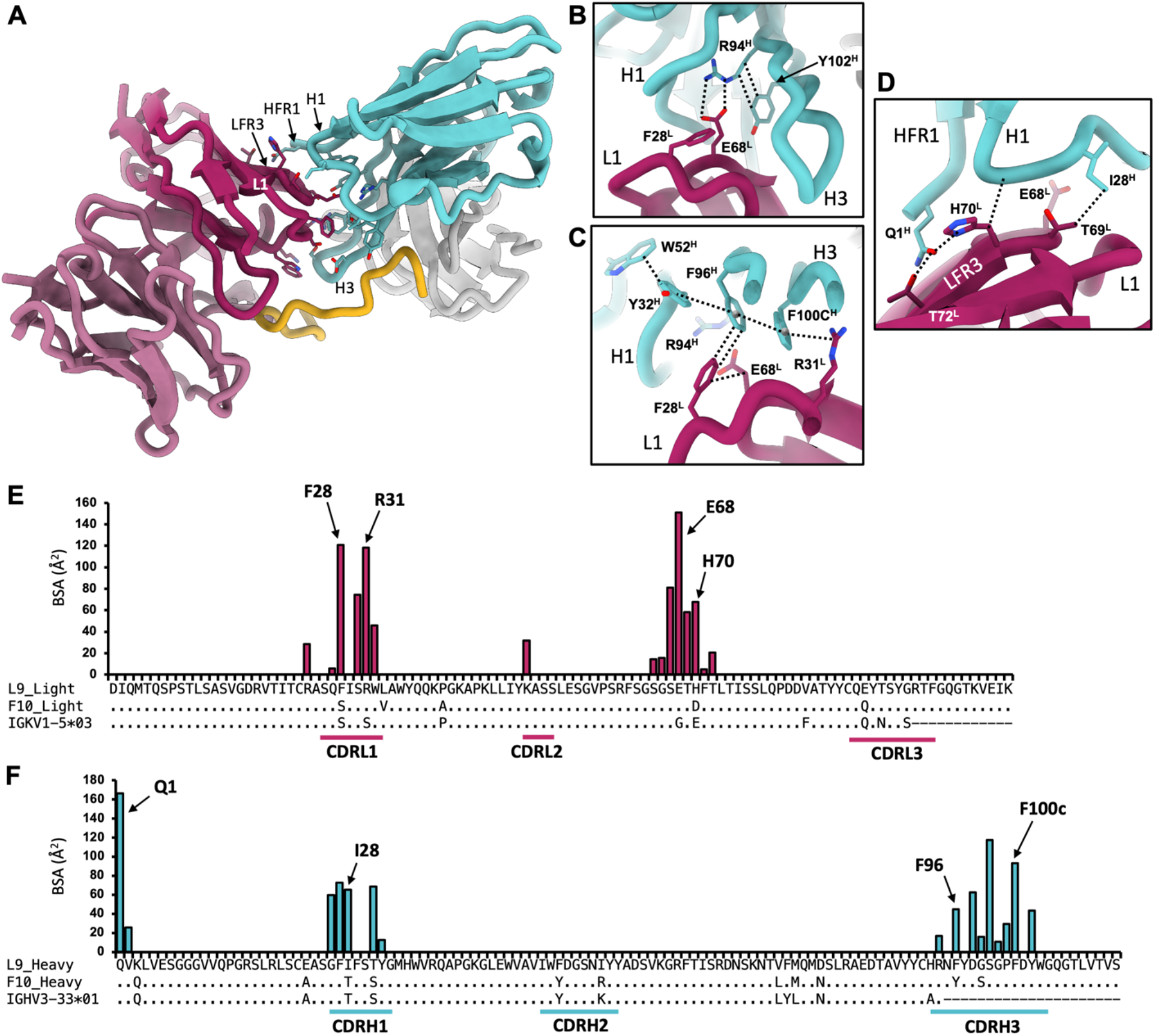
L9_K_ mediates extensive homotypic interactions. **(A)** Ribbon diagram of Fab B (cyan) and C (maroon); side chains of interacting residues are shown. **(B-D)** Structural details of key homotypic interactions. Dashed lines indicate specific contacts. **(E)** Buried surface area (BSA) contributions of individual residues to the homotypic interface in L9 light chain. Sequence alignment with F10_K_ and germline *IGKV1-5* gene is shown below. **(F)** Same as in (E), for L9 heavy chain, with sequence alignment to F10_H_ and germline *IGHV3-33* gene.

The L9 homotypic interface is distinct from that observed in NPNA-specific *IGHV3-33* mAbs, which is generally conserved and derives primarily from the heavy chain (*11*, *19*) (fig. S5). In contrast, L9_K_ contributes numerous critical homotypic contacts, and total BSA in the interface is evenly distributed between heavy and light chains (905Å^2^ and 839Å^2^, respectively) (Fig. 3, E and F). In the cryo-EM structure, L9_K_ of FabC packs tightly against L9_H_ of FabB, and extensive polar and hydrophobic contacts are made between CDRL1 and the LC framework region 3 of FabC (LFR3) with HFR1, CDRH1, and CDRH3 of FabB (Fig. 3A; fig. S4; table S3). The interface between FabB and FabA is nearly identical. Importantly, several residues mediating critical homotypic interactions (Fig. 3, B to D) correlate with somatic hypermutation of the germline *IGHV3-33* and *IGKV1-5* genes (Fig. 3E and F; fig. S5). Four somatically mutated residues in L9_K_, F28^L^ and R31^L^ in CDRL1, and E68^L^ and H70^L^ in LFR3, account for the majority of BSA contributed by the LC to the homotypic interface (Fig. 3E).

E68^L^ lies at the core of the homotypic interface in L9, where it forms a key salt bridge with the germline-encoded R94^H^ of CDRH3_B_ (Fig. 3B; fig. S4). In L9_H_, R94^H^ forms a conserved interaction with Y102^H^ to stabilize the base of CDRH3; thus E68^L^ may also indirectly impact antigen binding through stabilization of the CDRH3 loop in the adjacent Fab. F28^L^ coordinates a series of π-π stacking interactions in the opposing CDRH1_B_ (Y32^H^) and CDRH3_B_ (F96^H^ and F100c^H^) while also packing against the E68^L^ side chain. This pi network culminates in a cation-π bond between R31^L^ from CDRL1_C_ and F100c^H^ from the opposing CDRH3_B_ (Fig. 3C). On the other side of the homotypic interface from E68^L^, a mutated framework residue H70^L^ forms a hydrogen bond with the side chain of Q1^H^ in FabA in addition to multiple van der Waals contacts with CDRH1_B_ (Fig. 3D). Each of these homotypic contacts are not encoded in the germline sequence, and none directly contact rsCSP (fig. S3, A and B). These findings provide strong evidence for affinity maturation to optimize antibody-antibody binding, which may in turn enhance CSP avidity and protective efficacy, as we have shown recently for multiple NPNA-specific *IGHV3-33* mAbs (*11*).

The four somatic mutations in L9_K_ are atypical: F28^L^, E68^L^, and H70^L^ are observed in less than 1% percent of all human *IGKV1* light chain sequences, while R31^L^ is observed in only 2% (Fig. S5A) (*21*). Strikingly, F28 and H70 also correspond to two of the five amino-acid differences between mature L9 and the chimeric L9 mAb F10_K_/L9_H_ (S28 and D70 in F10_K_). As F28 and H70 both mediate key homotypic interactions in L9, which would likely be lost in F10_K_, these residues may explain the functional differences of F10_K_/L9_H_ from L9, namely (1) reduced avidity to CSP minor repeats, (2) loss of the ability to bind two adjacent NPNV repeats, and (3) significantly reduced protection *in vivo* (*p*<0.001) (*6*).

To test this hypothesis, and to understand the role of homotypic contacts in L9_K_ in general, we used molecular dynamics simulations to characterize WT L9 and a series of L9_K_ variants. L9_K_ residues were reverted to either the germline *IGKV1-5* gene (R31S, E68G) or to the L9_K_ precursor F10_K_ (F28S, H70D). We first compared the free energy landscapes of the CDR loops of individual Fv domains unbound to rsCSP (Fig. 4; fig. S6). We find that the R31S, E68G and H70D mutations in L9_K_ result in a broader conformational space and additional highly probable minima compared to the WT L9 Fv, indicating that these residues are critical for determining the shape and the conformational flexibility of the paratope (Fig. 4, B and C; fig. S6). These minima correspond to a substantial shift away from the binding competent conformation in combination with a higher conformational entropy, suggesting a decrease in stability and/or binding affinity (Fig. 4D). Importantly, when combined (R31S-E68G-H70D), these mutations significantly destabilize the homotypic interface (table S4; *p*<0.001), substantiating their key role in mediating homotypic interactions. Interestingly, the H70D single mutant *stabilizes* the homotypic interface (table S4), suggesting the germline E70 or F10_K_ D70 may have initialized the evolution of homotypic interactions during L9 maturation. Unlike other LC mutants, the F28S Fv reveals a similar conformational space and diversity in the CDR loops compared to the WT L9 Fv. However, F28S leads to formation of a new *intra*molecular salt bridge between residues R31^L^ and E68^L^, with simultaneous loss of the *inter*molecular salt bridge between E68^L^ and R94^H^ and the cation-π bond between R31^L^ and F100c^H^ (Fig. 4A). Thus, in addition to direct homotypic interactions, F28 acts indirectly through E68^L^ and R31^L^ to further stabilize antibody-antibody binding. This is reflected in the significantly decreased interaction energies of the homotypic interface in the F28S mutant relative to WT L9 (table S4) and is visualized in Movie S1. To understand the molecular basis of key functional differences between L9 and F10, we next modelled the F10 chimeras in the context of the trimeric Fab-rsCSP complex. Compared to WT L9 and L9_K_/F10_H_, the homotypic interface is strongly destabilized in F10_K_/L9_H_ (table S4). This suggests that F10_K_/L9_H_ would not bind multivalently to the minor repeats and would have overall reduced binding affinity, which is consistent with our previous functional data on this chimera (*6*). Five residues differ between L9_K_ and F10_K_: F28S, L33V, P40A, H70D, and E90Q (Fig. 3E). We find that the F28S mutation alone accounts for ~80% of the destabilization of the homotypic interface observed with F10_K_/L9_H_ compared to WT L9, while the H70D single mutant and the L33V-P40A-E90Q triple mutant Fvs both slightly *increase* stability of the complex (table S4). Taken together, these data suggest that the dramatic destabilization seen in MD simulations of the F10_K_/L9_H_ chimera is primarily the result of the F28S mutation. Therefore, this rare mutation in L9_K_ (S28F), and the network of homotypic contacts it mediates, may underlie the key functional differences between L9 and F10_K_/L9_H_.

**Figure 4.**
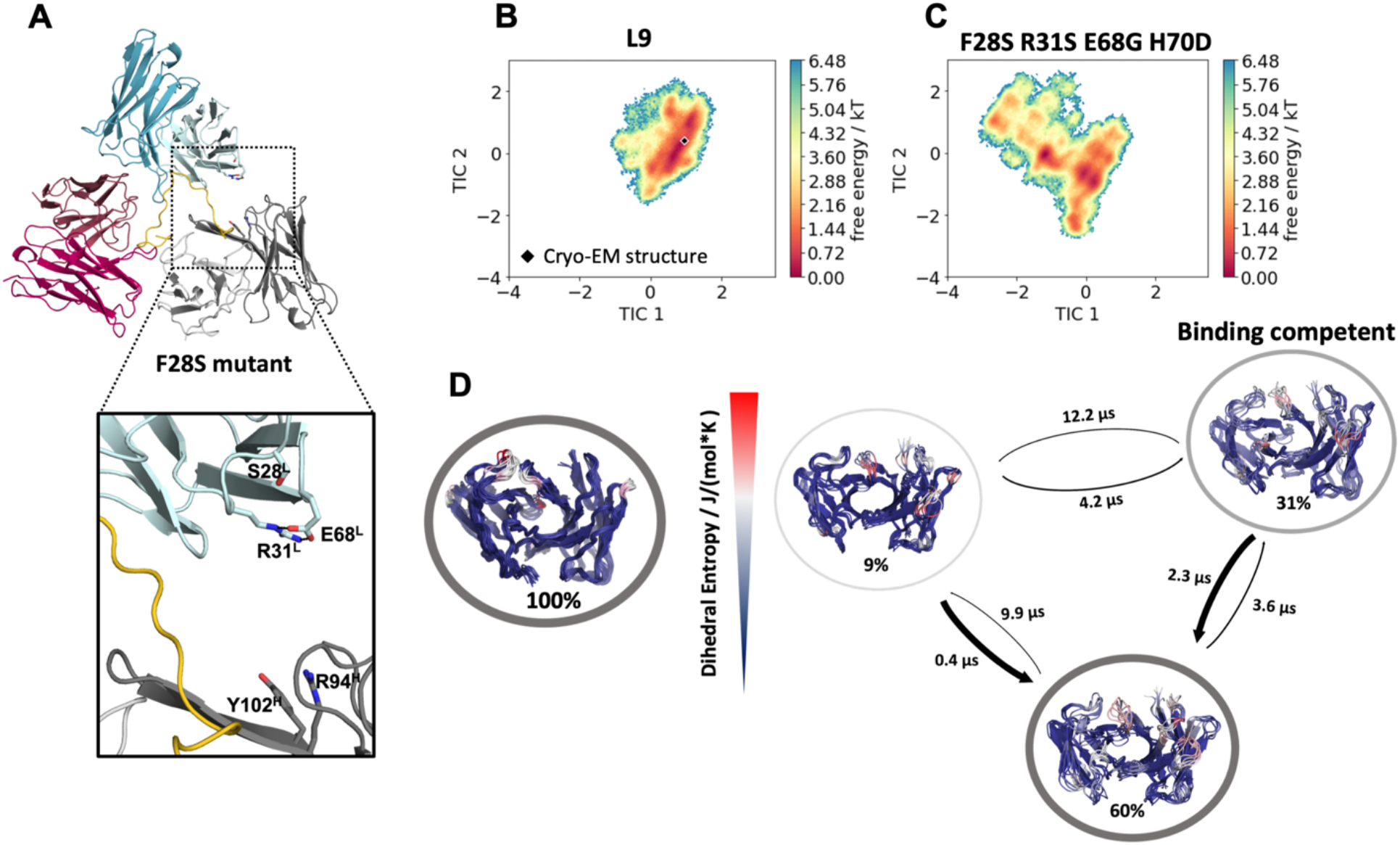
Molecular dynamics reveals L9_K_ residues critical for stability of the homotypic interface and CSP binding. **(A)** Most populated structure for the F28S variant, highlighting the loss of critical homotypic interactions. **(B-C)** Free energy landscapes of the L9 WT and the F28S/R31S/E68G/H70E variant projected in the same coordinate system, revealing a substantial increase in conformational space and a population shift due to the mutations. Cryo-EM structure is depicted as black diamond. **(D)** Conformational ensemble representatives, state probabilities and transition kinetics for the WT and the quadruple mutant, color-coded according to their dihedral entropy (blue-low flexibility, red-high variability).

Overall, this study reveals the structural basis for the extraordinary selectivity and binding affinity of L9 for the NPNV minor repeats and highlights the critical role of L9_K_ for both functions. We find that rare, somatically mutated residues in L9_K_ mediate extensive homotypic contacts between adjacent L9 Fabs and thus multivalent binding to adjacent NPNV motifs. These contacts underscore the requirement of at least two NPNV motifs for high affinity CSP binding by L9 (1000 nM vs 13 nM for CSP peptides with one and two NPNV, respectively) (*6*); Based on our recent finding that affinity-matured homotypic interactions in three potent NPNA-specific *IGHV3-33* mAbs are critical for both high NPNA avidity and protective efficacy (*11*), it is likely that L9_K_-mediated homotypic interactions are also critical for the potency of L9. Notably, these L9_K_ residues (F28, R31, E68, H70) make no direct contacts with rsCSP (fig. S3; table S2), indicating that the minor repeat region facilitates antibody-antibody affinity maturation in the context of multiple adjacent NPNV motifs, as has been observed for extended NPNA repeats (*11*, *18*, *19*). L9 is one of the most potent anti-PfCSP mAbs and is currently undergoing clinical development as a monoclonal therapy for malaria prevention (*17*). Thus, these structural data will be useful for rational antibody engineering to improve both the protective efficacy and pharmacokinetic properties of this mAb. The discovery of L9 and the NPNV minor repeat region as a highly protective epitope on PfCSP has led to new efforts to re-design PfCSP-based vaccines to elicit L9-like antibodies (*22*, *23*). The cryo-EM structure presented here now enables a structure-based approach, which may be instrumental in developing the next-generation malaria vaccine. Future studies to identify related, NPNV-specific mAbs should enhance our understanding of this class of antibodies and their important contribution to protective immunity against malaria.

## Acknowledgments

The authors thank B. Anderson for maintenance and administration of the cryo-EM facility at The Scripps Research Institute, and H.L. Turner and C.A. Bowman for technical support. We also thank L.T. Wang and N.K. Hurlburt for sharing of reagents and insightful discussions, and J.R. Riccabona and Y. Wang for fruitful discussions and technical support. The computational results presented here have been achieved (in part) using the Vienna Scientific Cluster (VSC). We acknowledge PRACE for awarding us access to Piz Daint at CSCS, Switzerland.

## Author contributions

GMM, MP, RAS, IAW, and ABW conceived the project. GMM, MFQ, WHL, and TP designed and performed experiments, and analyzed the data. LEW analyzed data. KRL, MP, RAS, IAW, and ABW acquired funding and supervised the project. GMM and MFQ wrote the original manuscript draft. All authors contributed to manuscript review and editing.

## Data and materials availability

The coordinates for the L9-rsCSP structure and the corresponding cryo-EM map have been deposited to the Protein Data Bank (PDB) and Electron Microscopy Data Bank (EMDB), respectively, with the accession codes 8EH5 and EMD-28135.

## Supplementary Materials

Materials and Methods

Figures S1-S4

Tables S1-S4

References 23-58

Movie S1

