## Supplemental Information for "Structural basis of epitope selectivity and potent protection from malaria by PfCSP antibody L9"


**This PDF file includes:**


Materials and Methods

Figures S1-S4

Tables S1-S4

References 24-60

Movie S1

#### MATERIALS AND METHODS

##### Protein production

L9 heavy and light chain sequences were codon-optimized for mammalian expression and cloned into pCMV. The recombinant Fab was expressed by transient transfection in Freestyle 239F cells (ThermoFisher), and purified with a Kappa affinity column followed by cation exchange chromatography (monS). rsCSP, which contained a C-terminal 6x-His tag, was cloned into pET28a and expressed in the Shuffle strain of *E. coli* (New England Biolabs). rsCSP was purified as previously described (*24*).

##### Cryo-EM sample preparation

To form the L9 Fab-rsCSP complex, an excess of L9 Fab was incubated with rsCSP in tris-buffered saline (TBS; pH 8.0) overnight at 4 °C. The complex was purified by size exclusion chromatography (SEC) with a Superdex 200 Increase 10/300 GL column (Sigma-Aldrich) equilibrated with TBS. For initial cryo-EM attempts, the purified complex was concentrated to ~1 mg/mL with a 30 kDa molecular weight cutoff filter (MilliporeSigma), and 3μL of this solution was applied to holey gold UltrAufoil (Quantifoil) cryo-EM grids. Grids were blotted for two to four seconds at 100% humidity, 4 °C, and plunge-frozen with a Vitrobot Mark IV into liquid ethane. Due to extensive aggregation of the complex during vitrification, and concomitant preferred orientation in vitrified ice, cryo-EM data were later collected with L9-rsCSP captured onto graphene oxide (GO) grids. For GO grid preparation, the L9-rsCSP complex was diluted to ~0.05 mg/mL in TBS, and 3 μL was applied to holey gold UltrAufoil grids containing a non-uniform layer of GO sheets on top of the grid. GO grids were made in-house, and fabrication preparation procedure was adapted from a published protocol (*25*). Briefly, UltrAufoil 1.2/1.3

holey gold grids (300 mesh) were washed with chloroform and allowed to dry completely. Grids were then glow-discharged, and 4 μL of 1 mg/mL PEI solution (polyethylenimine HCl, 25 mM HEPES pH 7.9) was applied to the grid and incubated for 2 minutes. Excess PEI was blotted with filter paper. Grids were washed with 2 drops of milli-Q water and allowed to dry completely. GO sheets (Sigma-Aldrich 763705) were diluted to 0.2 mg/mL in water and centrifuged at 1500xg. 4 μL of the supernatant was applied to grids and incubated for 2 minutes. Excess GO solution was blotted off, and grids were washed two times with water. Grids were allowed to dry for at least 30 minutes before use, and were used for sample vitrification on the same day they were prepared. This procedure resulted in ~90% coverage of the holes with GO; about half of these holes contained a monolayer of GO. For vitrification, 3 μL of L9-rsCSP complex (0.05 mg/mL) was applied to GO grids, and the sample was blotted for 2 sec at 100% humidity, 4 °C, and plunge-frozen in liquid ethane.

*Cryo-EM data collection*

Automated data acquisition was performed with the Leginon software (*26*) on a Titan Krios (ThermoFisher) operated at 300 keV. Micrograph movies were collected in electron counting mode with a K2 Summit direct electron detector (Gatan), with an unbinned pixel size of 1.045Å and a defocus range of -0.9μm to -2.0μm. The dose rate was ~6 $e^-/Å^2$/sec, with a full exposure time of 10 seconds; 200 ms per movie frame. This resulted in a total dose of ~60 $e^-/Å^2$ on the specimen. A total of 12,521 movies were collected over four separate data sets. Due to preferred orientation of the L9-rsCSP complex on GO grids, two of these four data sets were collected with a stage tilt of -40°, with all other imaging parameters held constant. Movies and micrographs were catalogued and stored with the aid of Appion (*27*).

*Single particle cryo-EM data processing*

Movie frames were aligned and dose-weighted with MotionCor2 (*28*). All subsequent processing was performed with cryoSPARCv3.3 (*29*). The contrast transfer function (CTF) was calculated with the Patch CTF Estimation tool, which was critical for accurate estimation of the tilted micrographs. The Gaussian (blob) picker was used on a subset of micrographs for initial particle picking, and 2D templates were generated with multiple rounds of 2D classification. Template picking was then used on the full dataset. Multiple rounds of 2D classification resulted in a particle stack containing 842,590 particles. A starting model generated from *ab initio* reconstruction was used for a non-uniform refinement job to achieve a resolution of ~3.7Å. Multiple rounds of global CTF (beam tilt) refinement and per particle defocus refinement led to a 3.35Å map. To account for possible flexibility between each of the three L9 Fabs, 3D Variability Analysis was used specifying 4 principal modes (*30*). The output was fed into a 3D Variability Display job in cluster mode, specifying 20 clusters. Close inspection of the interactive cluster plots and structural comparison of cluster maps identified rotational flexibility in the Fab (Fab A) bound at the N-terminus of the peptide relative to the other two Fabs. The most homogeneous clusters were pooled, yielding a particle stack with 451,712 particles. These were again subjected to non-uniform and CTF refinement, leading to a 3.36Å map with significantly improved interpretability of high-resolution features, particularly for the antigen (CSP) density.

*Atomic model building*

The X-ray structure of 239 Fab bound to $NPNA_2$ (6W00), which contains matching germline heavy chain (*IGHV3-33*) and light chain (*IGKV1-5*) genes, was used to generate a homology model of

L9 Fab. This model was then used as the template for re-building of the structure with RosettaCM (*31*). At first, only the central Fab was modelled. On the resulting lowest energy model, the CDR loops were removed and built manually in Coot (*32*). This structure was docked into the density of the two neighboring Fabs, and the trimeric Fab complex was refined with PHENIX real-space refine (*33*). Based on the known preferred epitope of L9, and inspection of the cryo-EM density, the structure of the PfCSP minor repeat region was built manually in Coot. The L9 Fab-rsCSP complex was again refined with PHENIX and errors were iteratively corrected with Coot. Rosetta Relax was used for a final all-atom refinement (*34*).

*Structural analysis*

Buried surface area (BSA) and root mean square deviation (RMSD) calculations were performed in UCSF Chimera (*35*). For general structural interpretation, UCSF Chimera and Coot were used. Calculation of electrostatic potential surfaces was performed with PyMol (The PyMOL Molecular Graphics System, Version 2.0 Schrödinger, LLC). The Epitope Analyzer webtool was used to assess direct contacts within the homotypic interface and between L9 Fab and rsCSP (tables S2 and S3; fig. S4) (*36*). Structure figures were made with UCSF Chimera, UCSF ChimeraX (*37*), and PyMol.

*Molecular dynamics simulations*

Based on the cryo-EM structure of the WT L9 (this study), containing three Fvs bound to rsCSP, we performed five replicas each of 1 μs of classical molecular dynamics simulations of the complex to identify critical residues that stabilize/favor the homotypic interface. For the other investigated variants (table S4), we derived the starting structures for our simulations from the WT

L9 structure by replacing the respective amino acids, followed by a local energy minimization in MOE (Molecular Operating Environment, Chemical Computing Group, version 2020.09). The starting structures for simulations were prepared in MOE using the Protonate3D tool (*38*). To neutralize the charges, we used the uniform background charge, which is required to compute long-range electrostatic interactions (*39*). Using the tleap tool of the AmberTools20 (*40*) package, the structures were soaked in cubic water boxes of TIP3P water molecules with a minimum wall distance of 12 Å to the protein (*41, 42*). For all simulations, parameters of the AMBER force field 14SB were used (*43*). Molecular dynamics simulations were performed in an NpT ensemble using pmemd.cuda (*44*). Bonds involving hydrogen atoms were restrained by applying the SHAKE algorithm (*45*), allowing a time step of 2 fs. Atmospheric pressure of the system was preserved by weak coupling to an external bath using the Berendsen algorithm (*46*). The Langevin thermostat was used to maintain the temperature during simulations at 300 K. The interaction energies were calculated with cpptraj by using the linear interaction energy (LIE) tool (*40*). We calculated the electrostatic and van der Waals interaction energies for all frames of each simulation (10000 frames/simulation) and provided the simulation-averages of these interaction energies in table S4.

A previously published method characterizing the CDR loop ensembles in solution (*47*) was used to investigate the conformational diversity of the six CDR loops of the free (apo) L9 Fv and the respective variants. To enhance the sampling of the conformational space, well-tempered bias-exchange metadynamics (*48, 49*) simulations were performed in GROMACS (*50, 51*) with the PLUMED 2 implementation (*52*). We chose metadynamics as it enhances sampling on predefined collective variables (CV). The sampling is accelerated by a history-dependent bias potential, which is constructed in the space of the CVs (*53*). As collective variables, we used a well-established protocol, boosting a linear combination of sine and cosine of the ψ torsion angles of all six CDR loops calculated with functions MATHEVAL and COMBINE implemented in PLUMED 2 (*47*). As discussed previously, the ψ torsion angle captures conformational transitions comprehensively (*54*). The underlying method presented in this paper has been validated in various studies against a large number of experimental results (*47, 55*). The simulations were performed at 300 K in an NpT ensemble using the GPU implementation of the pmemd module (*44*) to be as close to the experimental conditions as possible and to obtain the correct density distributions of both protein and water. We used a Gaussian height of 10.0 kJ/mol and a width of 0.3 rad. Gaussian deposition occurred every 1000 steps and a biasfactor of 10 was used. 500 ns of bias-exchange metadynamics simulations were performed for the prepared Fv structures. The resulting trajectories were aligned to the whole Fv and clustered with cpptraj (*40*) using the average linkage hierarchical clustering algorithm with a RMSD cut-off criterion of 1.2 Å resulting in a large number of clusters. The cluster representatives for the antibody fragments were equilibrated and simulated for 100 ns using the AMBER 20 simulation package. The accumulated simulation times for the investigated L9 variants are summarized in table S5.

With the obtained trajectories, we performed a time-lagged independent component analysis (tICA) using the python library PyEMMA 2 employing a lag time of 10 ns. tICA was applied to identify the slowest movements of the investigated Fv fragments and consequently to obtain a kinetic discretization of the sampled conformational space (*56*). tICA is a dimensionality reduction technique that detects the slowest-relaxing degrees of freedom and facilitates kinetic clustering, which is a crucial pre-requisite for building a Markov-state model. It linearly transforms a set of high-dimensional input coordinates to a set of output coordinates, by finding a subspace of *"good* *reaction coordinates"*. Thereby, tICA finds coordinates of maximal autocorrelation at a given lag time. The lag time sets a lower limit to the timescales considered in the tICA and the Markov-state model. Accordingly, tIC1 and tIC2 represent the two slowest degrees of freedom of the systems.

Based on the tICA conformational spaces, thermodynamics and kinetics were calculated with a Markov-state model (MSM) (*57*) by using PyEMMA 2, which uses the k-means clustering algorithm to define microstates and the PCCA+ clustering algorithm (*58*) to coarse-grain the microstates to macrostates. Markov-state models are network models which provide valuable insights for conformational states and transition probabilities between them, as it allows identification of the boundaries between two states (*57*). Basically, MSMs coarse-grain the system's dynamics, which reflect the free energy surface and ultimately determine the system's structure and dynamics. Thus, MSMs provide important insights and enhance the understanding of states and transition probabilities and facilitates a quantitative connection with experimental data (*59*).

The sampling efficiency and the reliability of the Markov-state model (e.g., defining optimal feature mappings) has been evaluated with the Chapman-Kolmogorov test by using the variational approach for Markov processes and monitoring the fraction of states used, since the network states must be fully connected to calculate probabilities of transitions and the relative equilibrium probabilities. To build the Markov-state model, we used the backbone torsions of the respective CDR loops, defined 100 microstates using the k-means clustering algorithm and applied a lag time of 10 ns.

Additionally, we calculated the residue-wise dihedral entropies with the recently published X-entropy python package, which calculates the entropy of a given dihedral angle distribution (*60*). This approach uses a Gaussian kernel density estimation (KDE) with a plug-in bandwidth selection, which is fully implemented in C++ and parallelized with OpenMP. The obtained residue-wise dihedral entropies were projected onto the respective structures (Figure 4D).

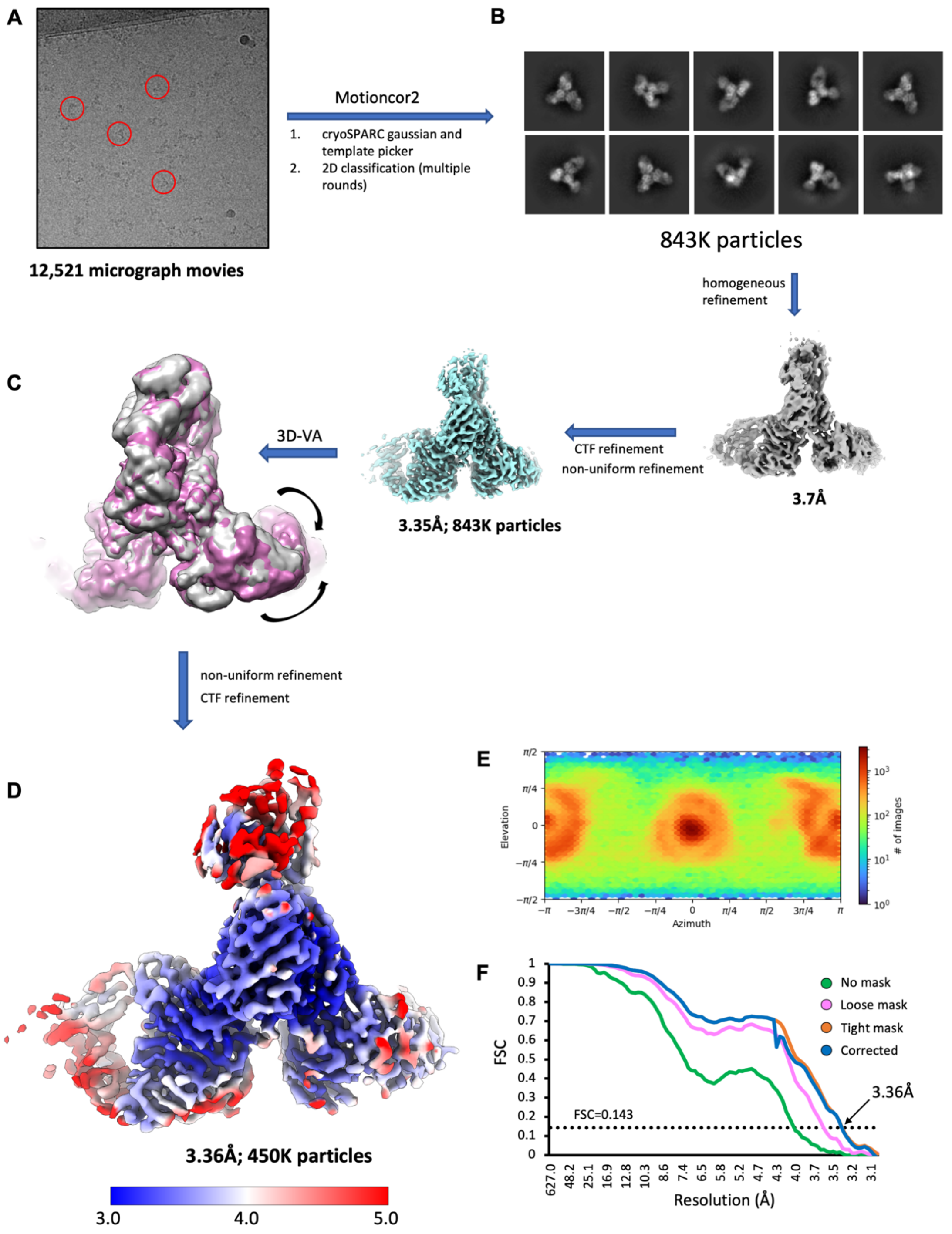

A
12,521 micrograph movies
Motioncor2
1. cryoSPARC gaussian and template picker
2. 2D classification (multiple rounds)
B
843K particles
homogeneous refinement
3.7Å
CTF refinement
non-uniform refinement
3.35Å; 843K particles
3D-VA
C
non-uniform refinement
CTF refinement
D
3.36Å; 450K particles
3.0
4.0
5.0
E
Elevation
Azimuth
### of images
F
FSC
Resolution (Å)
No mask
Loose mask
Tight mask
Corrected
FSC=0.143
3.36Å

**Figure S1. Cryo-EM reconstruction of L9-rsCSP and data processing workflow. (A)** Representative cryo-EM micrograph of L9 on graphene oxide. **(B)** Representative 2D class averages. **(C)** Intermediate cryo-EM maps. Right: initial consensus refinement; Middle: high-resolution map with full data set; Right: representative cluster maps from 3D variability analysis (VA) of full dataset, showing motion in Fab A. **(D)** Final map after 3D-VA and further CTF refinement. **(E)** Angular distribution plot of final map in (D). **(F)** Fourier shell correlation (FSC) plot of final map.

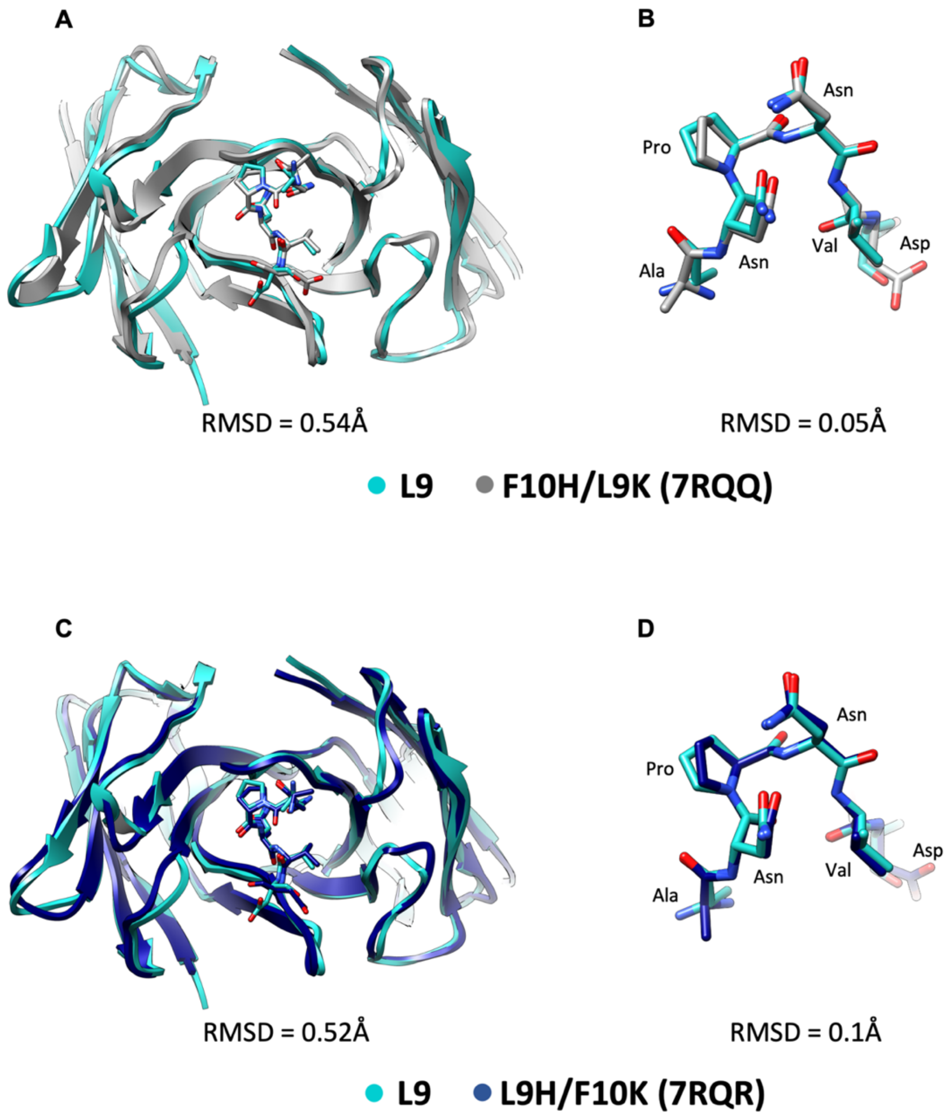


**Figure S2. Structural comparison of L9 Fab cryo-EM structure with X-ray structures of L9 chimeras. (A)** Overlay of single L9 Fab and rsCSP (ANPNVD) with F10$_H$/L9$_K$ bound to NANPNVD, where NPNV forms a type 1 β-turn. **(B)** Structural match of CSP peptides from L9 cryo-EM and F10$_H$/L9$_K$ X-ray structures. **(C,D)** Same as in (A) and (B), for L9$_H$/F10$_K$.

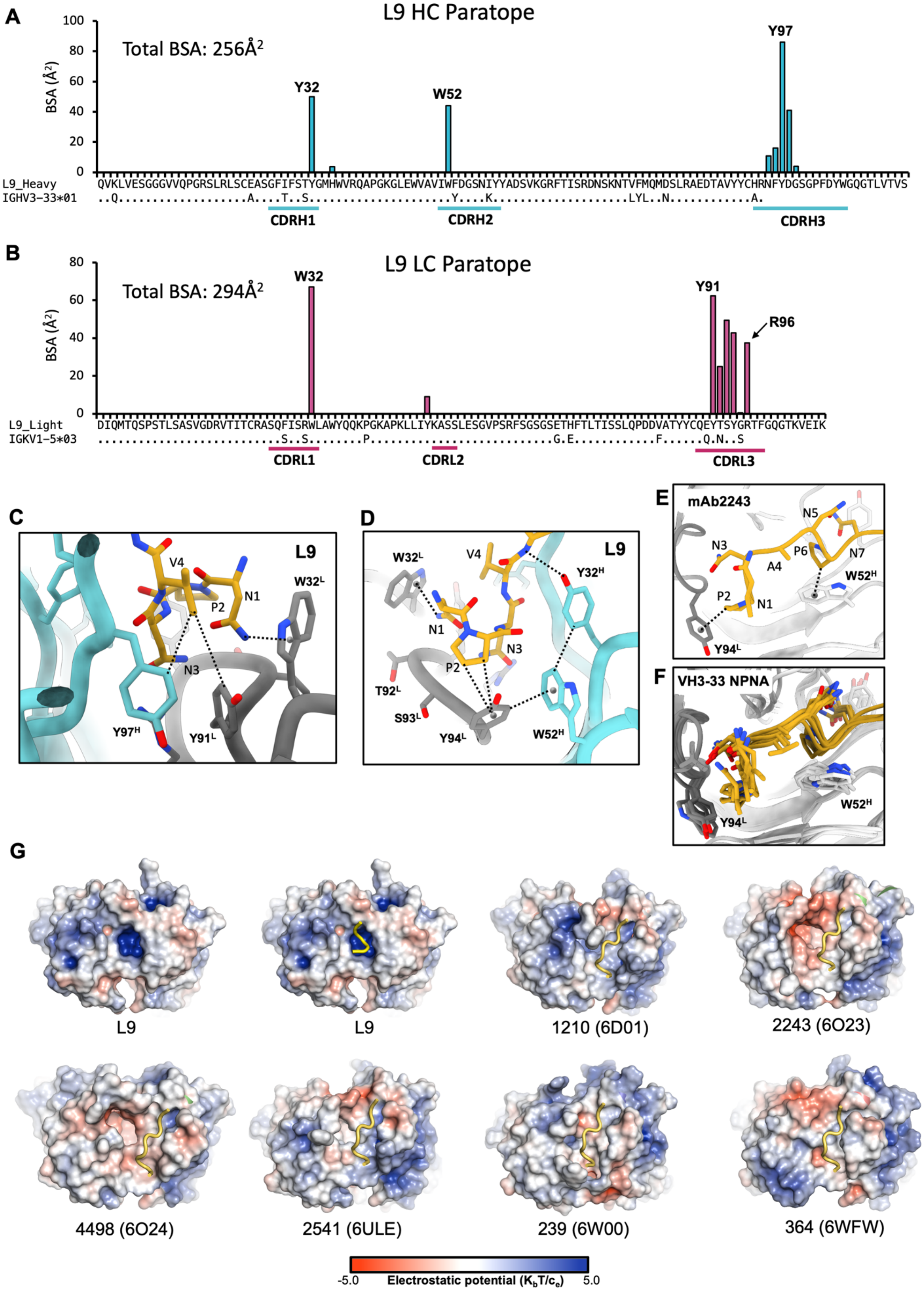

A
L9 HC Paratope
Total BSA: 256Å²
BSA (Å²)
100
80
60
40
20
0
Y32
W52
Y97
L9_Heavy QVKLVESGGGVVQPGRSLRLSCEASGFIFSTYGMHWVRQAPGKGLEWVAVIWFDGSNIYYADSVKGRFTISRDNSKNTVFMQMDSLRAEDTAVYYCHRNFYDGSGPFDYWGQGTLVTVS
IGHV3-33*01 ..Q....................A....T..S...................Y....K..................LYL..N...........A.
CDRH1
CDRH2
CDRH3
B
L9 LC Paratope
Total BSA: 294Å²
BSA (Å²)
80
60
40
20
0
W32
Y91
R96
L9_Light DIQMTQSPSTLSASVGDRVTITCRASQFISRWLAWYQQKPGKAPKLLIYKASSLESGVPSRFSGSGSETHFTLTISSLQPDDVATYYCQEYTSYGRTFGQGTKVEIK
IGKV1-5*03 ...........................S..S.......P..........................G.E...........F......Q.N..S
CDRL1
CDRL2
CDRL3
C
L9
V4
P2
N1
N3
W32ᴸ
Y97ᴴ
Y91ᴸ
D
L9
W32ᴸ
V4
Y32ᴴ
N1
N3
P2
T92ᴸ
S93ᴸ
Y94ᴸ
W52ᴴ
E
mAb2243
N3
N5
P6
N7
A4
P2
N1
W52ᴴ
Y94ᴸ
F
VH3-33 NPNA
W52ᴴ
Y94ᴸ
G
L9
L9
1210 (6D01)
2243 (6O23)
4498 (6O24)
2541 (6ULE)
239 (6W00)
364 (6WFW)
-5.0
Electrostatic potential (K_bT/c_e)
5.0

**Figure S3. Structural details of the L9 paratope.** **(A)** Buried surface area (BSA) contributions of individual residues to CSP binding in the L9 heavy chain. Sequence alignment to the germline *IGHV3-33* germline gene shown below. **(B)** Same as in (A), for the L9 light chain. **(C,D)** Structural details of NPNV binding. **(E)** $NPNA_2$ epitope structure in the NPNA-specific mAb 2243 (PBD 6O23). **(F)** Same as in (E), with X-ray structures of six NPNA-specific mAbs superimposed to highlight structural conservation. These six mAbs are shown in (G). **(G)** Electrostatic surface potentials from L9 cryo-EM structure +/- peptide (upper left two panels) and X-ray structures of six other NPNA-specific mAbs bound to peptide. The PDB accession codes are in parentheses.

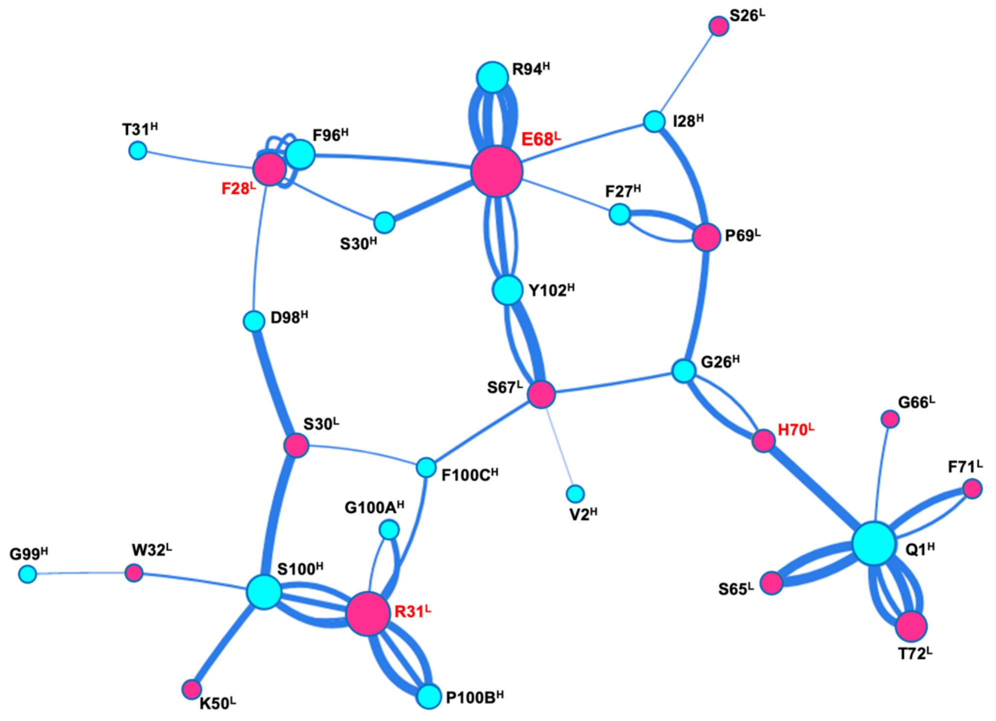


**Figure S4. Contact network of the L9 homotypic interface.** Light chain residues are colored in magenta; heavy chain residues are in cyan. The size of each circle corresponds to the overall contribution of that residue to the homotypic interface. The width of the lines indicates the strength of the homotypic contact. Generated with the Epitope Analyzer webtool (*36*).

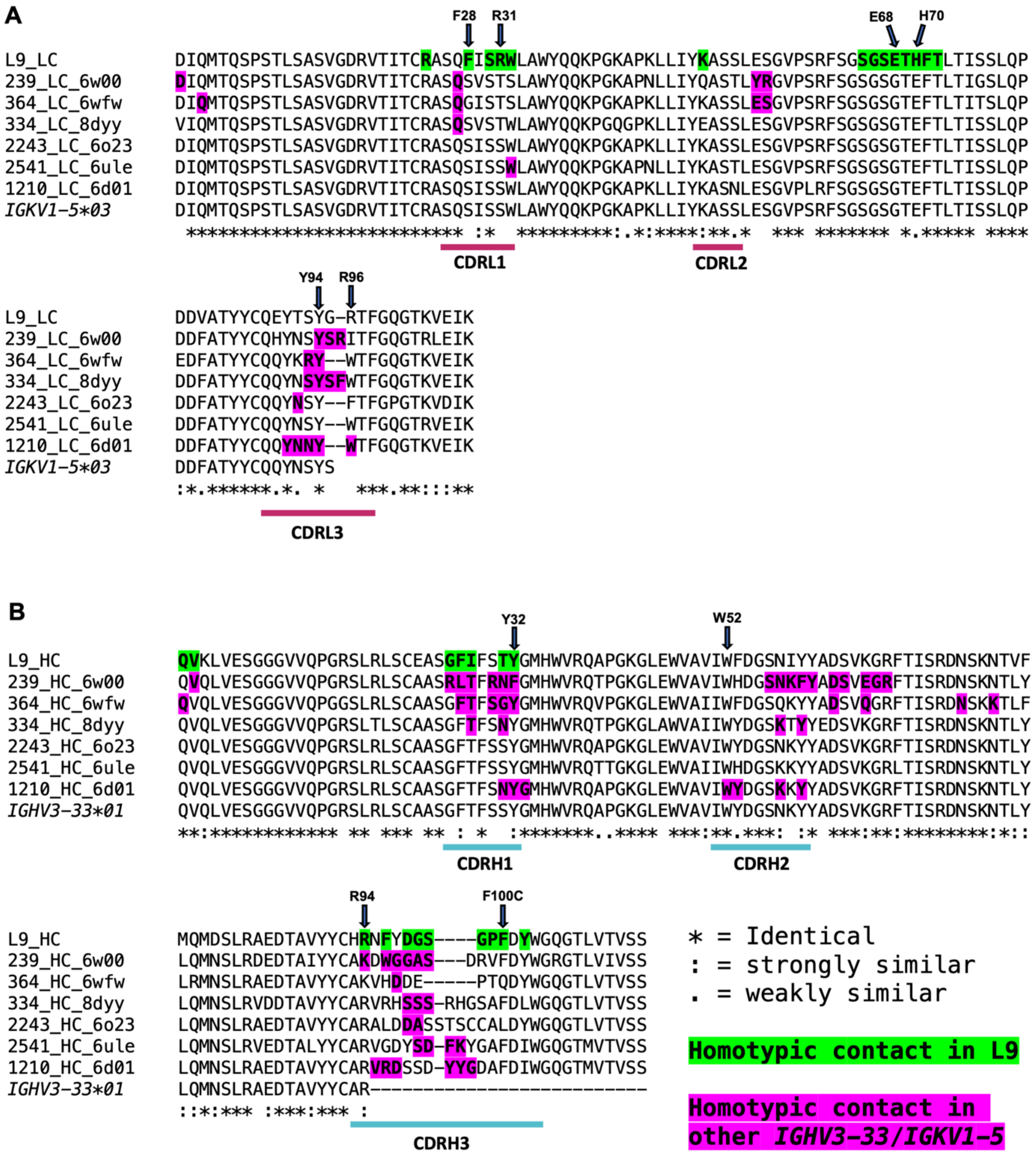

**Figure S5.** Multiple sequence alignment (MSA) of L9 with NPNA-targeting mAbs all utilizing *IGKV1-5* and *IGHV3-33* light and heavy chains, respectively. PDB codes follow mAb names. **(A)** Light chain MSA. **(B)** Heavy chain MSA.

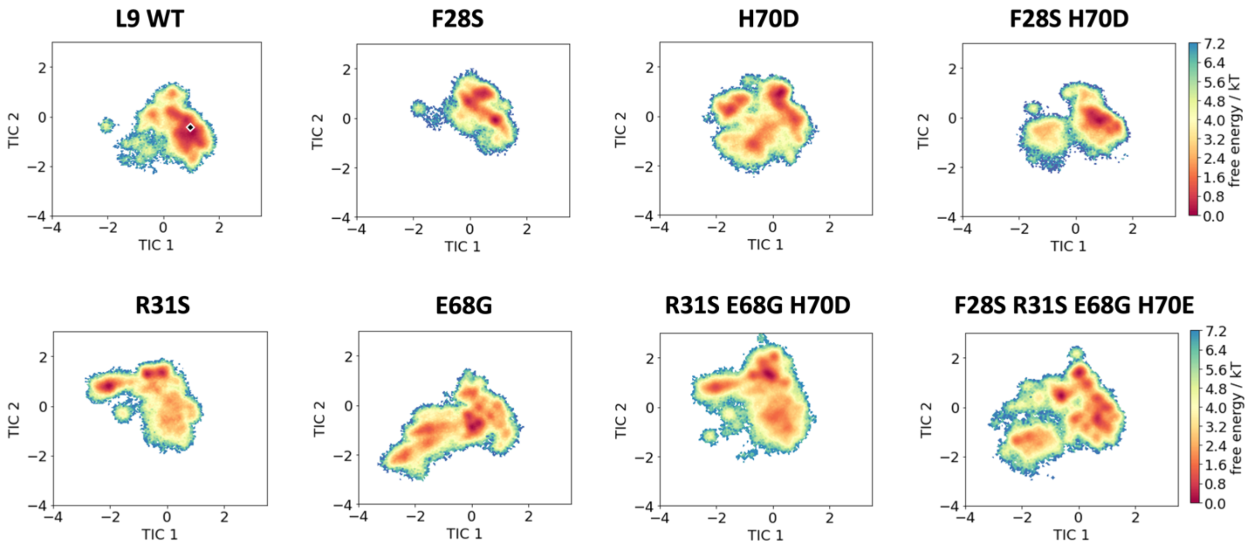


**Figure S6.** Free energy landscapes of isolated Fv domains of all variants modelled. See also Figure 4.

**Table S1.** Cryo-EM data collection parameters and model statistics

| | L9 Fab-rsCSP<br>PDB: 8EH5<br>EMDB: 28135 |
|---|---|
| **Data Collection** | |
| Microscope | Titan Krios |
| Detector | Gatan K2 Summit |
| Voltage (kV) | 300 |
| Pixel Size (Å) | 1.045 |
| Defocus range (µm) | -0.9 to -2.0 |
| Total electron dose ($e^-/Å^2$) | 60 |
| Dose rate ($e^-/Å^2$/sec) | 6 |
| Frames per exposure | 50 |
| **Data Processing** | |
| Total micrograph movies | 12,521 |
| Particle images in map | 451,712 |
| Symmetry imposed | C1 |
| Map resolution (FSC=0.143; Å) | 3.36 |
| Map sharpening B-factor ($Å^2$) | 136.9 |
| Data processing software | cryoSPARC v3.3 |
| **Model Refinement** | |
| No. atoms in deposited model | 5509 |
| Chains total | 7 |
| Residues (protein) | 704 |
| RMS Deviations | |
| Bond lengths (Å) | 0.022 |
| Bond angles (°) | 1.719 |
| Validation | |
| MolProbity score | 1.37 |
| Clash score | 2.69 |
| EMRinger score | 3.22 |
| Poor rotamers (%) | 0.5 |
| Ramachandran plot | |
| Favored (%) | 95.5 |
| Allowed (%) | 3.9 |
| Outliers (%) | 0.6 |
| Average B-factor | 47 |

**Table S2.** Direct contacts between central L9 Fab and rsCSP

| Chain 1 | Residue 1 | Position 1 | Atom (1>2) | Chain 2 | Residue 2 | Position 2 | Distance (Å) | Predicted interaction |
|---|---|---|---|---|---|---|---|---|
| G | ASN | 115 | ND2-CD2 | L | TRP | 32 | 3.2 | Van-der-Waals |
| G | ASN | 115 | ND2-O | L | TYR | 91 | 2.9 | H-bond |
| G | ASN | 115 | ND2-CA | L | THR | 92 | 3.7 | Van-der-Waals |
| G | ASN | 115 | ND2-O | L | THR | 92 | 3.9 | H-bond |
| G | PRO | 116 | O-CE2 | H | TYR | 32 | 3.4 | Van-der-Waals |
| G | PRO | 116 | O-CZ2 | H | TRP | 52 | 3.6 | Van-der-Waals |
| G | PRO | 116 | CD-O | L | THR | 92 | 3.9 | Van-der-Waals |
| G | PRO | 116 | CD-CA | L | SER | 93 | 4.3 | Van-der-Waals |
| G | PRO | 116 | CB-CZ | L | TYR | 94 | 3.6 | Hydrophobic |
| G | PRO | 116 | CB-CE2 | L | TYR | 94 | 3.6 | Hydrophobic |
| G | PRO | 116 | CG-CZ | L | TYR | 94 | 3.7 | Hydrophobic |
| G | PRO | 116 | CG-CE1 | L | TYR | 94 | 3.8 | Hydrophobic |
| G | PRO | 116 | CB-CE1 | L | TYR | 94 | 4.1 | Hydrophobic |
| G | PRO | 116 | CG-CE2 | L | TYR | 94 | 4.2 | Hydrophobic |
| G | PRO | 116 | CB-CD2 | L | TYR | 94 | 4.3 | Hydrophobic |
| G | PRO | 116 | CG-CD1 | L | TYR | 94 | 4.3 | Hydrophobic |
| G | ASN | 117 | OD1-CH2 | H | TRP | 52 | 4.0 | Van-der-Waals |
| G | ASN | 117 | OD1-O | H | ASN | 95 | 4.1 | Van-der-Waals |
| G | ASN | 117 | O-CA | H | PHE | 96 | 3.4 | Van-der-Waals |
| G | ASN | 117 | O-N | H | TYR | 97 | 2.9 | H-bond |
| G | ASN | 117 | ND2-O | L | TYR | 91 | 3.5 | H-bond |
| G | ASN | 117 | ND2-O | L | SER | 93 | 2.9 | H-bond |
| G | ASN | 117 | OD1-NH2 | L | ARG | 96 | 2.8 | H-bond |
| G | ASN | 117 | OD1-NH1 | L | ARG | 96 | 2.9 | H-bond |
| G | VAL | 118 | C-OH | H | TYR | 32 | 4.1 | Van-der-Waals |
| G | VAL | 118 | O-OH | H | TYR | 32 | 4.4 | H-bond |
| G | VAL | 118 | CA-O | H | TYR | 97 | 3.5 | Van-der-Waals |
| G | VAL | 118 | CG2-CB | H | TYR | 97 | 4.2 | Hydrophobic |
| G | VAL | 118 | CG2-CD2 | H | TYR | 97 | 4.3 | Hydrophobic |
| G | VAL | 118 | N-O | H | TYR | 97 | 4.4 | H-bond |
| G | VAL | 118 | CG1-CB | H | TYR | 97 | 4.4 | Hydrophobic |
| G | VAL | 118 | CG2-CE1 | L | TYR | 91 | 4.3 | Hydrophobic |
| G | ASP | 119 | CB-OH | H | TYR | 32 | 3.4 | Van-der-Waals |
| G | ASP | 119 | N-OH | H | TYR | 32 | 3.6 | H-bond |
| G | ASP | 119 | OD2-OH | H | TYR | 32 | 4.4 | H-bond |
| G | ASP | 119 | OD1-OH | H | TYR | 32 | 4.4 | H-bond |
| G | ASP | 119 | N-O | H | TYR | 97 | 3.2 | H-bond |
| G | ASP | 119 | CB-OD1 | H | ASP | 98 | 3.4 | Van-der-Waals |

**Table S3.** Homotypic interactions in L9-rsCSP cryo-EM structure (Fab B and Fab C)

| Chain Fab 1 | Residue 1 | Position 1 | Atom (1>2) | Chain Fab 2 | Residue 2 | Position 2 | Distance (Å) | Predicted interaction |
|---|---|---|---|---|---|---|---|---|
| P (L) | SER | 26 | O-CD1 | H | ILE | 28 | 4.1 | Van-der-Waals |
| P (L) | PHE | 28 | CB-OG | H | SER | 30 | 3.8 | Van-der-Waals |
| P (L) | PHE | 28 | CE1-OG1 | H | THR | 31 | 4.0 | Van-der-Waals |
| P (L) | PHE | 28 | CZ-CE1 | H | PHE | 96 | 3.5 | Hydrophobic |
| P (L) | PHE | 28 | CE1-CE1 | H | PHE | 96 | 3.5 | Hydrophobic |
| P (L) | PHE | 28 | CE1-CZ | H | PHE | 96 | 3.8 | Hydrophobic |
| P (L) | PHE | 28 | CZ-CD1 | H | PHE | 96 | 4.2 | Hydrophobic |
| P (L) | PHE | 28 | CZ-CZ | H | PHE | 96 | 4.2 | Hydrophobic |
| P (L) | PHE | 28 | CE1-CD1 | H | PHE | 96 | 4.5 | Hydrophobic |
| P (L) | PHE | 28 | CE2-OD2 | H | ASP | 98 | 3.9 | Van-der-Waals |
| P (L) | SER | 30 | OG-OD2 | H | ASP | 98 | 2.8 | H-bond |
| P (L) | SER | 30 | O-OG | H | SER | 100 | 2.8 | H-bond |
| P (L) | SER | 30 | CA-CZ | H | PHE | 100C | 4.1 | Van-der-Waals |
| P (L) | ARG | 31 | CD-OG | H | SER | 100 | 3.6 | Van-der-Waals |
| P (L) | ARG | 31 | NH1-O | H | SER | 100 | 3.6 | H-bond |
| P (L) | ARG | 31 | NH1-OG | H | SER | 100 | 4.2 | H-bond |
| P (L) | ARG | 31 | NE-OG | H | SER | 100 | 4.3 | H-bond |
| P (L) | ARG | 31 | NH1-CA | H | GLY | 100A | 3.8 | Van-der-Waals |
| P (L) | ARG | 31 | NH1-O | H | GLY | 100A | 4.4 | H-bond |
| P (L) | ARG | 31 | NH2-O | H | PRO | 100B | 2.8 | H-bond |
| P (L) | ARG | 31 | NH1-O | H | PRO | 100B | 3.5 | H-bond |
| P (L) | ARG | 31 | NH1-N | H | PRO | 100B | 4.1 | H-bond |
| P (L) | ARG | 31 | NE-CE2 | H | PHE | 100C | 3.3 | Van-der-Waals |
| P (L) | TRP | 32 | CZ2-C | H | GLY | 99 | 4.2 | Van-der-Waals |
| P (L) | TRP | 32 | CZ2-N | H | SER | 100 | 3.7 | Van-der-Waals |
| P (L) | LYS | 50 | NZ-O | H | SER | 100 | 3.8 | H-bond |
| P (L) | SER | 65 | O-NE2 | H | GLN | 1 | 2.6 | H-bond |
| P (L) | SER | 65 | OG-NE2 | H | GLN | 1 | 2.9 | H-bond |
| P (L) | GLY | 66 | N-NE2 | H | GLN | 1 | 3.8 | Van-der-Waals |
| P (L) | SER | 67 | CB-CG2 | H | VAL | 2 | 4.4 | Van-der-Waals |
| P (L) | SER | 67 | O-O | H | GLY | 26 | 3.5 | Van-der-Waals |
| P (L) | SER | 67 | OG-CD2 | H | PHE | 100C | 3.4 | Van-der-Waals |
| P (L) | SER | 67 | OG-OH | H | TYR | 102 | 2.2 | H-bond |
| P (L) | SER | 67 | O-OH | H | TYR | 102 | 4.5 | H-bond |
| P (L) | GLU | 68 | OE1-CB | H | PHE | 27 | 3.9 | Van-der-Waals |
| P (L) | GLU | 68 | OE1-O | H | ILE | 28 | 3.6 | Van-der-Waals |
| P (L) | GLU | 68 | OE1-OG | H | SER | 30 | 4.1 | H-bond |
| P (L) | GLU | 68 | OE2-NE | H | ARG | 94 | 2.8 | H-bond |
| P (L) | GLU | 68 | OE2-NH2 | H | ARG | 94 | 3.6 | Salt-Bridge |
| P (L) | GLU | 68 | OE1-NH2 | H | ARG | 94 | 3.8 | Salt-Bridge |
| P (L) | GLU | 68 | OE1-NE | H | ARG | 94 | 4.1 | H-bond |
| P (L) | GLU | 68 | OE2-CZ | H | PHE | 96 | 3.1 | Van-der-Waals |
| P (L) | GLU | 68 | OE2-CE1 | H | TYR | 102 | 3.2 | Van-der-Waals |
| P (L) | GLU | 68 | OE2-OH | H | TYR | 102 | 3.9 | H-bond |
| P (L) | GLU | 68 | N-OH | H | TYR | 102 | 4.5 | H-bond |
| P (L) | THR | 69 | OG1-O | H | GLY | 26 | 3.9 | H-bond |
| P (L) | THR | 69 | OG1-CA | H | PHE | 27 | 3.5 | Van-der-Waals |
| P (L) | THR | 69 | OG1-N | H | PHE | 27 | 4.1 | H-bond |
| P (L) | THR | 69 | OG1-N | H | ILE | 28 | 3.8 | H-bond |
| P (L) | HIS | 70 | ND1-OE1 | H | GLN | 1 | 2.8 | H-bond |
| P (L) | HIS | 70 | CB-O | H | GLY | 26 | 3.4 | Van-der-Waals |
| P (L) | HIS | 70 | N-O | H | GLY | 26 | 3.9 | H-bond |
| P (L) | PHE | 71 | C-OE1 | H | GLN | 1 | 3.4 | Van-der-Waals |
| P (L) | PHE | 71 | N-OE1 | H | GLN | 1 | 3.7 | H-bond |
| P (L) | THR | 72 | OG1-OE1 | H | GLN | 1 | 2.9 | H-bond |
| P (L) | THR | 72 | N-OE1 | H | GLN | 1 | 3.4 | H-bond |
| P (L) | THR | 72 | OG1-NE2 | H | GLN | 1 | 3.6 | H-bond |
| P (L) | THR | 72 | O-NE2 | H | GLN | 1 | 4.4 | H-bond |

**Table S4. Interaction energies of homotypic interface in trimeric L9 Fv-rsCSP complexes.**

| L9 Variant | Electrostatic Energy (kcal/mol) | SD | van der Waals Energy (kcal/mol) | SD | Sim. Time (µs) |
|---|---|---|---|---|---|
| L9 | -150.2 | 32 | -27.4 | 6 | 5 |
| F28S* | -46.8 | 34 | -9.5 | 5 | 5 |
| F28S, H70D* | -82.6 | 28 | -16.4 | 6 | 5 |
| H70D* | -228.3 | 27 | -41.4 | 8 | 5 |
| E68G* | -39.3 | 44 | -11.8 | 6 | 5 |
| R31S | -170.4 | 30 | -29.9 | 5 | 5 |
| R31S, E68G, H70D* | -41.8 | 31 | -14.7 | 8 | 5 |
| F28S, R31S, E68G, H70D* | -18.9 | 33 | -6.1 | 6 | 5 |
| $F10_K$ $L9_H$ Chimera* | -20.2 | 32 | -5.6 | 6 | 5 |
| $L9_K$ $F10_H$ Chimera | -118.4 | 37 | -12.3 | 7 | 5 |
| L33V, P40A, E90Q | -170.8 | 31 | -23 | 6 | 5 |

* $p$<0.001; T-test, relative to L9

**Table S5. Free Fv (apo) aggregated simulation times.**

| L9 Variant | Aggregated simulation time (µs) |
|---|---|
| L9 WT | 21.6 |
| F28S | 18.5 |
| F28S, H70D | 19.3 |
| H70D | 23.4 |
| R31S | 15.9 |
| E68G | 19.1 |
| R31S, E68G, H70D | 23.9 |
| F28S, R31S, E68G, H70D | 25.8 |
